# Green Conversations: Harnessing Plant Communication to control growth light intensity

**DOI:** 10.1101/2024.08.29.610229

**Authors:** James Stevens, Phillip Davey, Piotr Kasznicki, Tanja A Hofmann, Tracy Lawson

**Affiliations:** School of Life Sciences, University of Essex, Colchester CO4 3SQ, UK

**Keywords:** basil, chlorophyll fluorescence, LEDs, plant communication, plant feedback system, photosynthetic efficiency, photosynthesis energy use, resource use, yield gain

## Abstract

Controlled Environment Agriculture (CEA) delivers increased crop production per unit land, contributing to resilient food systems amidst challenges of climate change, population growth and urbanization. However, high energy costs associated with lighting impose substantial barriers to the widespread adoption of CEA. While light is indispensable for growth, critically its utilization by crops throughout the photoperiod remains sub-optimal, reducing photosynthetic efficiency and wasting energy. Here we have developed and demonstrated a novel real-time plant bio-feedback system that enables crops to directly ‘communicate’ optimal lighting requirements. Continuous non- invasive monitoring of photochemistry elicited decreased demand for light by basil at the end of the photoperiod. Our innovative approach increased yield by 10% and reduced energy consumption per unit fresh mass by 18%, delivering a 201gCO_2_ gFW^-1^ reduction in carbon footprint. Application of this technique at scale can revolutionise resource management of CEA, reinvigorating the productivity, profitability and sustainability of this food industry.

**Author Contributions:** JS & TL: designed the experiments; JS, PD and PK: performed all physiology experiments and data acquisition and carried out data analyses. TL, JS, PD and TH wrote the MS and all authors commented on the MS.

**One-sentence summary:** Chlorophyll fluorescence measurements of photosynthetic efficiency can be used to control growth light intensity in real time, optimising crop performance and energy use.

## Introduction

Sustainable increases in food production are significant political and social challenges. Key demographic drivers include the growing global population^1^ and rising urbanization^2^. Meanwhile unpredictable weather patterns resulting from climate change add to those pressures^3^. As of 2020, agriculture itself contributes 14% of global CO2e emissions^4^, while yield gains are plateauing^5^; both issues should be addressed urgently if global net zero targets and food security goals are to be realised.

Controlled Environment Agriculture (CEA) combines scientific knowledge with technological advancements to address the pressures on the sector^6^. CEA systems rely on cultivating crops within partially or entirely enclosed environments, which give growers a significant degree of control over growing conditions^7,8^. Horticulturalists are already able to reduce water, nutrient and pesticide inputs in CEA while increasing productivity^9^. However, high electricity prices pressurise the financial viability of CEA where energy may represent up to 30% of operating costs^10^. Yet the industry holds great promise for the production of local, nutritious food while reducing transport costs and environmental impacts^11^.

Vertical Farms (VFs) are entirely closed, controlled environments where the grower must supply all inputs including lights^12^. Increased cropping area per unit land can be coupled with year-round production capabilities^13^. Those crops can be cultivated with enhanced nutritional value^14^. These factors fuel support for the VF industry, particularly in regions such as the Middle East and South East Asia^6,15^.

VFs are acutely sensitive to energy costs^16^, making environmental sustainability and the capacity of the industry to achieve net zero questionable^17^. Nevertheless CEA systems could lower CO emissions to the same level as broadacre agriculture by exploiting renewable energy sources^18^.

Elsewhere, progress on emission reductions is most likely to come from technological advancements^19^. Here we present a solution based on the design and development of a photo- feedback system that allows the crop to “communicate” directly with the LED growth lights, boosting productivity and promoting sustainability.

Currently the majority of indoor growth environments utilize a ‘square wave’ light regime, which maintains light intensity at a constant level over the photoperiod^20,21,22,23^. Light intensity is closely correlated to photosynthetic rate (*A*), which drives crop growth, directly linking lighting conditions to final yield^24^.

However, there is a significant drop in *A* at high light intensities and towards the end of the photoperiod^25,26^. Thus light use efficiency is not always directly proportional to intensity, even under square wave conditions^27^. Therefore, there is potential for substantial energy savings by dynamically matching light intensity to physiological demand^23^.To be commercially applicable, such an approach would require non-invasive measurements of photosynthetic performance^12^.

Typically, measurements of photosynthetic carbon assimilation using infra-red gas analysis are labour intensive and time-consuming^28^, limiting attractiveness to industry. An alternative technique is to measure *chlorophyll fluorescence* (CF), a rapid, scaleable and non-invasive method to determine the efficiency of PSII photochemistry (*Fq’*/*Fm’*), a proxy for *A* ^29,30^. Chlorophyll fluorescence is linked directly to photosynthesis by the rate of electron transport (ETR), in which the light energy absorbed by pigments (including chlorophylls) drives the flow of electrons through photosystems II (PSII) and I (PSI). ATP and a reductant (NADPH) are produced and used to fix CO_2_ in the Calvin-Benson-Bassham cycle to produce sugars and other biological molecules for growth^31^. However, two further processes compete for absorbed energy. In the first case, energy is dissipated as heat, known as *non-photochemical quenching* (NPQ)^29^. Alternatively, photons are re-emitted at a longer wavelength^29^. This latter process is chlorophyll fluorescence. As light intensity increases, photosynthetic efficiency (*Fq’/Fm’*) falls, and NPQ is a component of this decrease. Light intensities beyond saturation can result in damage to the PS’s, known as photoinhibition, further decreasing *Fq’/Fm’* (Baker, 2008).

Additionally, factors other than light, such as temperature or nutrients, can also affect *Fq’/Fm’*. Consequently, CF is often employed to assess plant health in a broad spectrum of conditions^32,33^.

Several CEA studies have shown improved edible biomass of indoor crops by lowering light intensity and increasing photoperiod (compared to higher light intensities for a shorter duration)^34,35,36,37^.

However, these studies failed to take into account the decrease in photosynthetic efficiency over the diurnal period.

Van Iersel at al.^38,39^ developed a biofeedback system that linked plant photosynthetic efficiency to light intensity via CF over a single photoperiod. They demonstrated that it was possible to maintain a wide range of ETRs and to distinguish between NPQ and photoinhibition to explain any reductions in photosynthetic efficiency^38,39^. Whilst these studies provided proof of concept, they did not assess the impact of such a feedback system on crop growth/yield and energy efficiency. Integration of real-time measurements of CF with precision control of LEDs represents a major technological advancement. This offers an exciting opportunity to adjust light based on the real-time physiological responses of the plant over the entire crop cycle^40,41^.

By adjusting light intensity to match photosynthetic demand, we demonstrated a photo-feedback system that enhanced basil yield while *<u>simultaneously</u>* reducing energy consumption. The result was a substantial improvement in cost effectiveness of CEA systems. By dynamically adjusting LED intensity based on photosynthetic efficiency during the growing period, we highlight the immense potential in engaging in “green conversations”.

## Results

Supporting observations in other species^26,27,42,43^, basil grown under conventional lighting regimes exhibited a substantial 15% shortfall in potential carbon gain (Fig. 1A). Maximum photosynthesis (*A*) of approximately 14 µmol m^-2^ s^-1^ was only sustained for the first 6 h of this 18 h square wave light regime. Subsequently, *A* steadily decreased to approximately 10 µmol m^-2^ s^-1^ by the end of the photoperiod.

**Figure 1.**
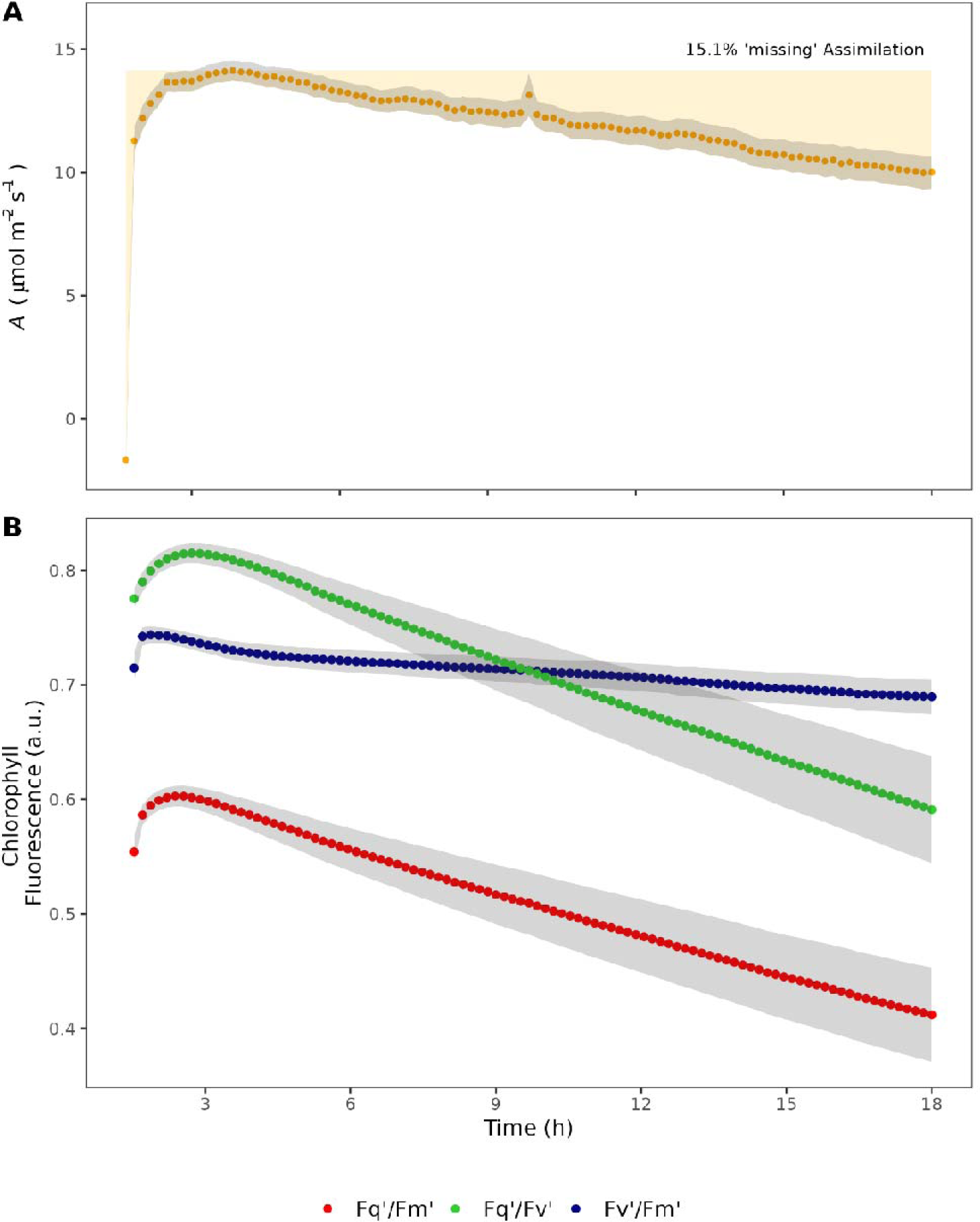
Basil (O. basilicum cv Sweet Genovese) plant responses to constant light conditions. Plants were exposed to 1000 µmol m^-2^ s^-1^ PAR for 18 h at 24° C, 410 ppm CO_2_ & VPD 1.8 kPa. A) Photosynthetic CO_2_ assimilation (A) showed a rise on exposure to light that continued for several h to peak between 3 h and 6 h. Thereafter assimilation fell steadily for the remainder of the photoperiod.B) Fq’/Fm’, the operating efficiency of Photosystem II (PSII) is a function of Fq’/Fv’ (the PSII efficiency factor) and Fv’/Fm’ (maximal quantum efficiency of PSII). Changes in Fq’/Fm’ were largely the result of changes in Fq’/Fv’ implying that a falling efficiency factor drove the decrease in operating efficiency. Fv’/Fm’ exhibited minimal change over the course of the measurement. Means shown +/- SE, n=6.

Photosynthetic efficiency is a function of maximum operating efficiency at the prevailing light intensity (*Fv’/Fm’*) and the proportion open reaction centres (*Fq’/Fv’*). The observed decline in *A* was driven by a reduction in the operating efficiency of photosystem II (PSII, *Fq’/Fm’*) (Fig. 1B).

Mechanistically this was caused by a decreased sink for the products of electron transport (*Fq’/Fv’*). Maximal capacity to transfer absorbed light energy to the photosystems (*Fv’/Fm’*) remained unaltered (Fig. 1B).

The close coupling between *A* and *Fq’/Fm’*allowed us to exploit rapid, robust and cost-effective chlorophyll fluorescence measurements to develop a real time photo-feedback system (Fig. 2). The relationship between light intensity and photosynthetic carbon assimilation (Fig. 3A) followed the expected hyperbolic response. The response of photosynthetic efficiency to light intensity was modelled as exponential decay (Fig. 3B). These responses were used to determine system light intensity limits and the setpoint (Fig 3. and Materials & Methods section). A proportional integral (PI) feedback loop managed light intensity around a setpoint within the capabilities of commercial LED lighting systems.

**Figure 2.**
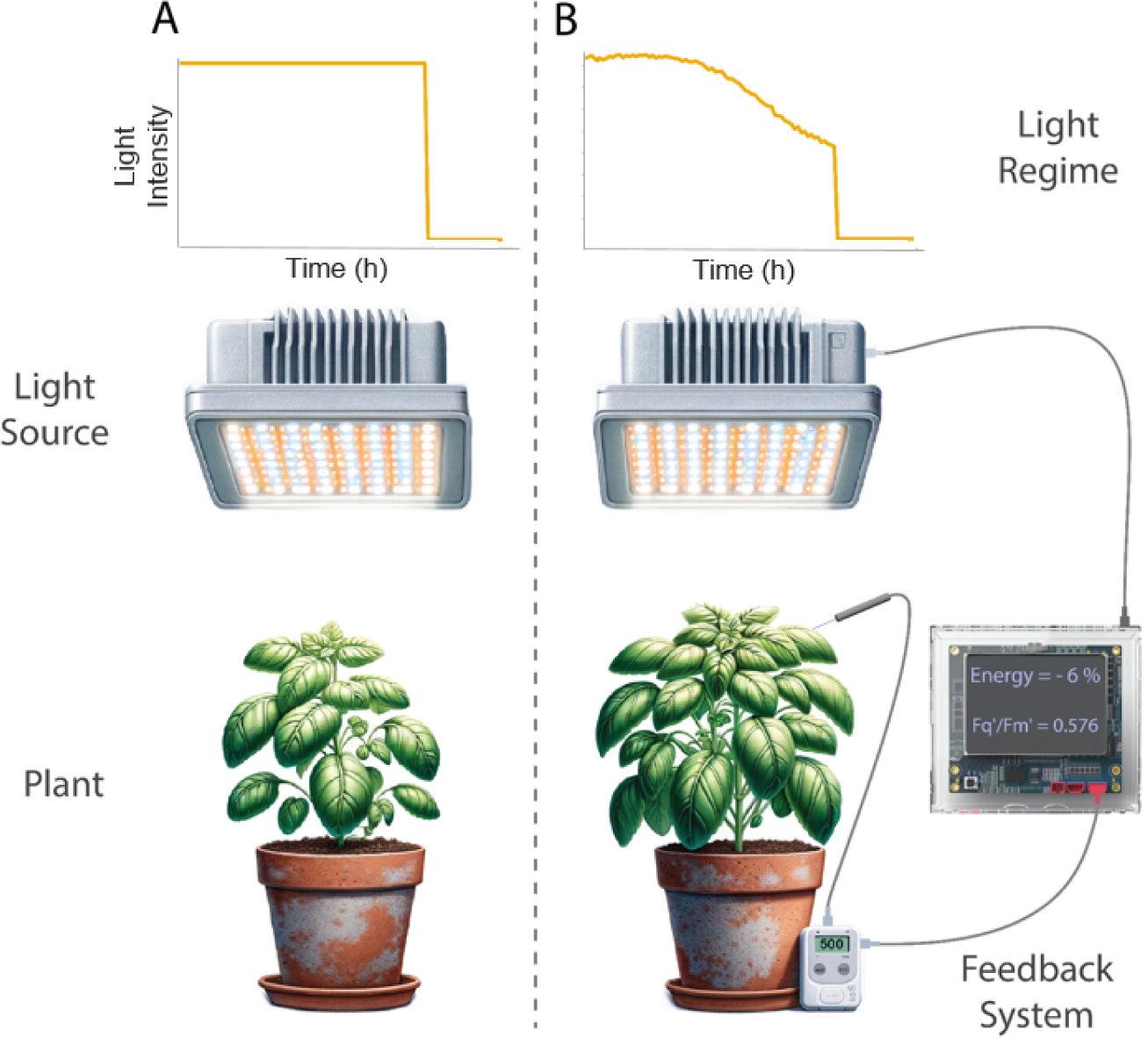
After seedling stage, basil plants were placed in one of two growth conditions for the duration of the experiment as soon as the youngest fully expanded leaves were large enough to be measured effectively. A) In the ’Controlled’ treatment, plants were illuminated in a temperature-controlled environment of 23°C for 18 h in 24 at 380 µmol m^-2^ s^-1.^ B) In the ’Feedback’ treatment, plants were probed using a OptiSciences PSP32 Chlorophyll fluorescence probe and datalogger. Variance in Fq’ / Fm’ compared to an empirically-defined setpoint of 0.52 was passed through a Proportion-Integration control loop and corrective intensity of illumination applied by a connection to a Heliospectra Dyna LED luminaire in a close loop. Illumination in the Feedback system was limited to a maximum of 450 µmol m^-2^ s^-1^ and a minimum of 100 µmol m^-2^ s^-1^. Day length was identical to the Control at 18 h.

**Figure 3.**
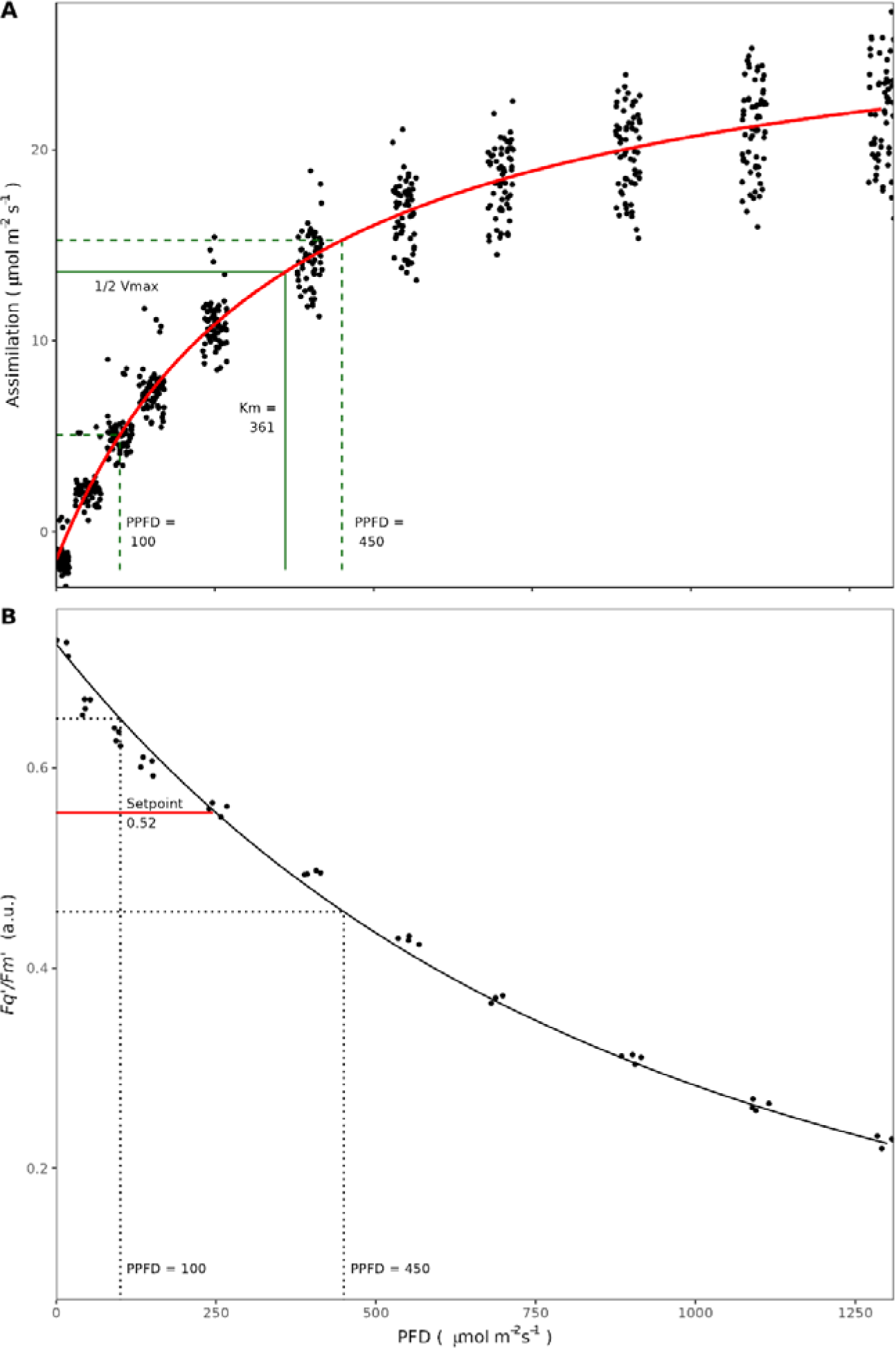
Key feedback parameters for the operation of the ’Green Conversation’ Chlorophyll Fluorescence device were an empirical process. A). Photosynthetic CO_2_ assimilation (A) as a function of light intensity (black points) was modelled as a Michaelis-Menten enzyme kinetic (red line). The inflection point ½V_max_, where the plants were most sensitive to a change in light intensity, was observed at a value K_m_ = 361 µmol m^-2^ s^-1^ PPFD. K_m_ lay between the minimum and maximum selected values of 100 µmol m^-2^ s^-1^ and 450 µmol m^-2^ s^-1^ which represent the economically viable range of intensities used by growers. n=20. B). The response of the operating efficiency of photosystem II as a function of PFD was modelled as a Gomperz function. The position of the empirically determined setpoint of Fq’/Fm’ at 0.52 lay at approximately the midpoint of Fq’/Fm’ observed at the maximum and minimum light intensity settings. n=6.

Our photo-feedback system produced a basil crop with substantial increases in key commercial yield parameters. Fresh and dry biomass and leaf area were significantly higher (p<0.01, p<0.001 and p<0.001 respectively, Fig. 4A-C). Harvestable fresh mass, a critical determinant of retail value, increased by 7.4 %, equating to 0.9 g per plant.

**Figure 4.**
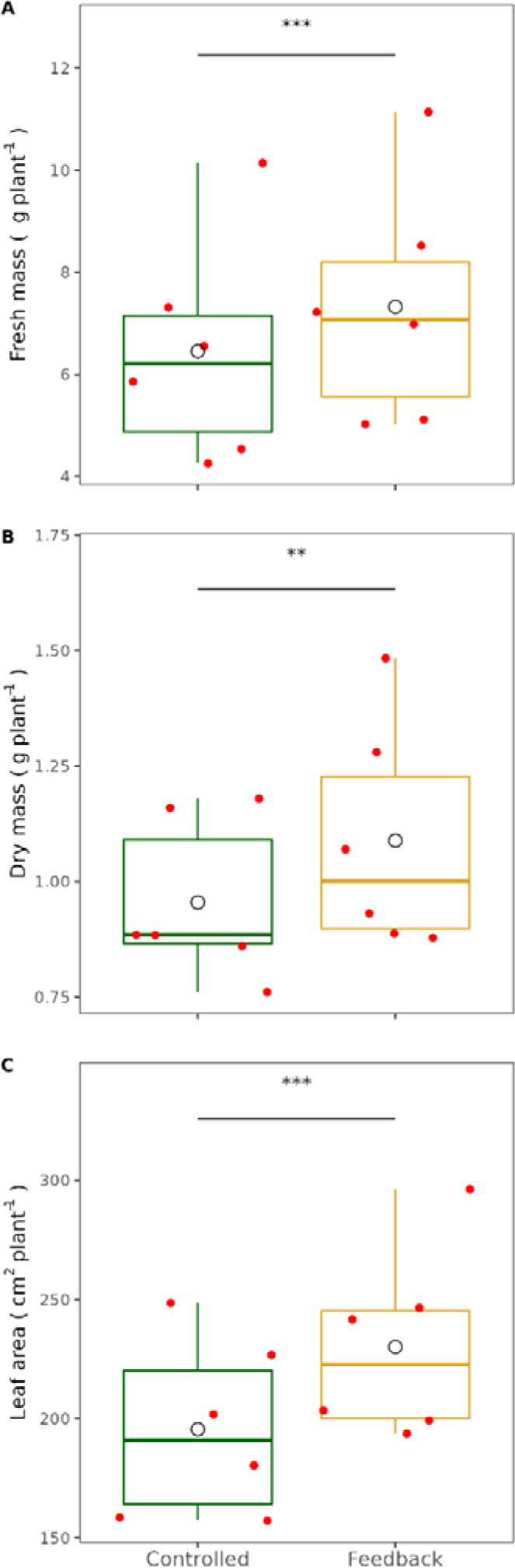
The harvest results of the Feedback system consistently outperformed the Controlled- lighting system. Data were analysed using a linear mixed effects model of the form lme(Harvest ∼ Treatment + (1|Experiment_number)). This approach captured more of the uncertainty in the (semi-controlled) environmental conditions experienced compared to a traditional linear model. Aikike Information Criteria were consistent for all results, with the mixed-effects model outperforming the linear model. A) Fresh mass of above-ground basil plants were significantly higher in the Feedback condition, B) Dry mass of above-ground basil plants were also significantly higher in the Feedback condition, C) Meanwhile, mean leaf area of basil plants were significantly higher in the Feedback condition. Red points represent individual experiments and the black circle is the overall mean for each treatment, n=6. Significance levels:*** = 0.001, ** = 0.01, * = 0.05.

Importantly these higher yields were achieved at a significantly lower total energy demand for lighting (p<0.01, Fig. 5A), representing a mean saving of 1.55 kWh or 6.18%, and resulting in significantly higher mean fresh mass, dry mass and leaf area *per unit energy consumed* (Fig. 5B-D; p<0.001). Once again, the commercially important metric of fresh mass yield per unit power increased by 20.2%. Moreover, growth using the photo-feedback system increased production of foliar area by 1.96 cm^2^ kWh^-1^ or 25.0% per plant. Crucially, no significant differences in relative water content or specific leaf area were observed (Supplementary data).

**Figure 5.**
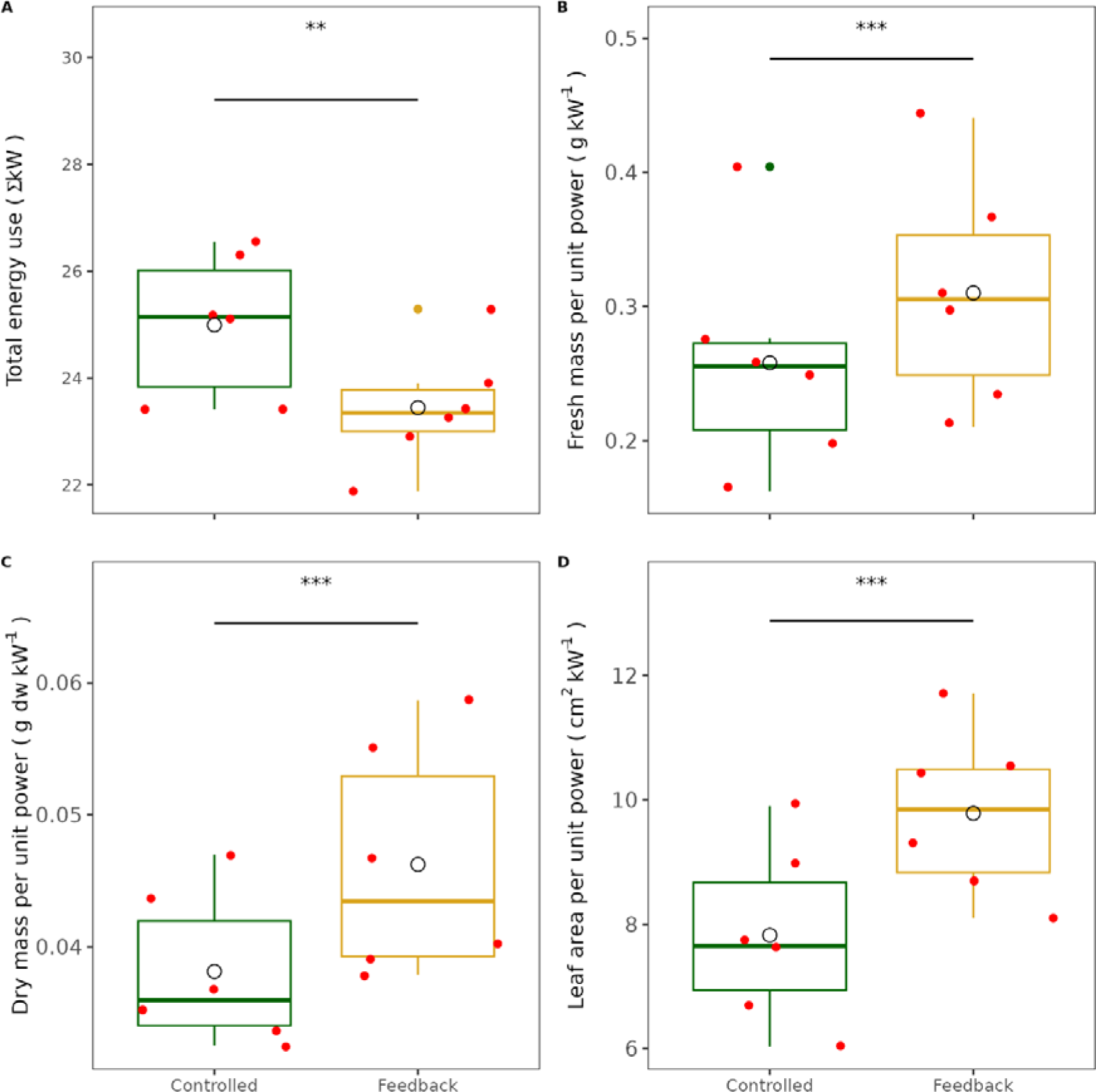
Power use was reduced in the Feedback condition. A) Total power use in each experiment was significantly lower in the Feedback condition. B) Fresh mass achieved per kW was significantly higher in the Feedback condition, as was C) dry mass per kW and D) Leaf area per kW. Red points represent individual experiments, black points show outliers and the black circle is the overall mean for each treatment, n=6. Significance levels:*** = 0.001, ** = 0.01, * = 0.05.

The photo-feedback system achieved gains in yield *and* savings in energy use by optimising light intensity over the diurnal period tuned to photosynthetic demand. Despite a significantly higher photosynthetic ‘demand’ during the first 9 h (e.g. at 3 h, p<0.01), light requirement subsequently dropped below the square wave light regime’s for the remainder of the photoperiod (e.g. at 15 h, p<0.001) (Fig 6A). There were no differences in *Fq’/Fm’* in the initial 9 h between the control and photo-feedback system (Fig. 6B). The real-time monitoring of chlorophyll fluorescence enabled the system to fine-tune light intensity and maintain *Fq’ / Fm’* close to the target of 0.52. This adjustment prevented the steep decline in operating efficiency seen in the control after 9 h (Fig. 6B).

**Figure 6.**
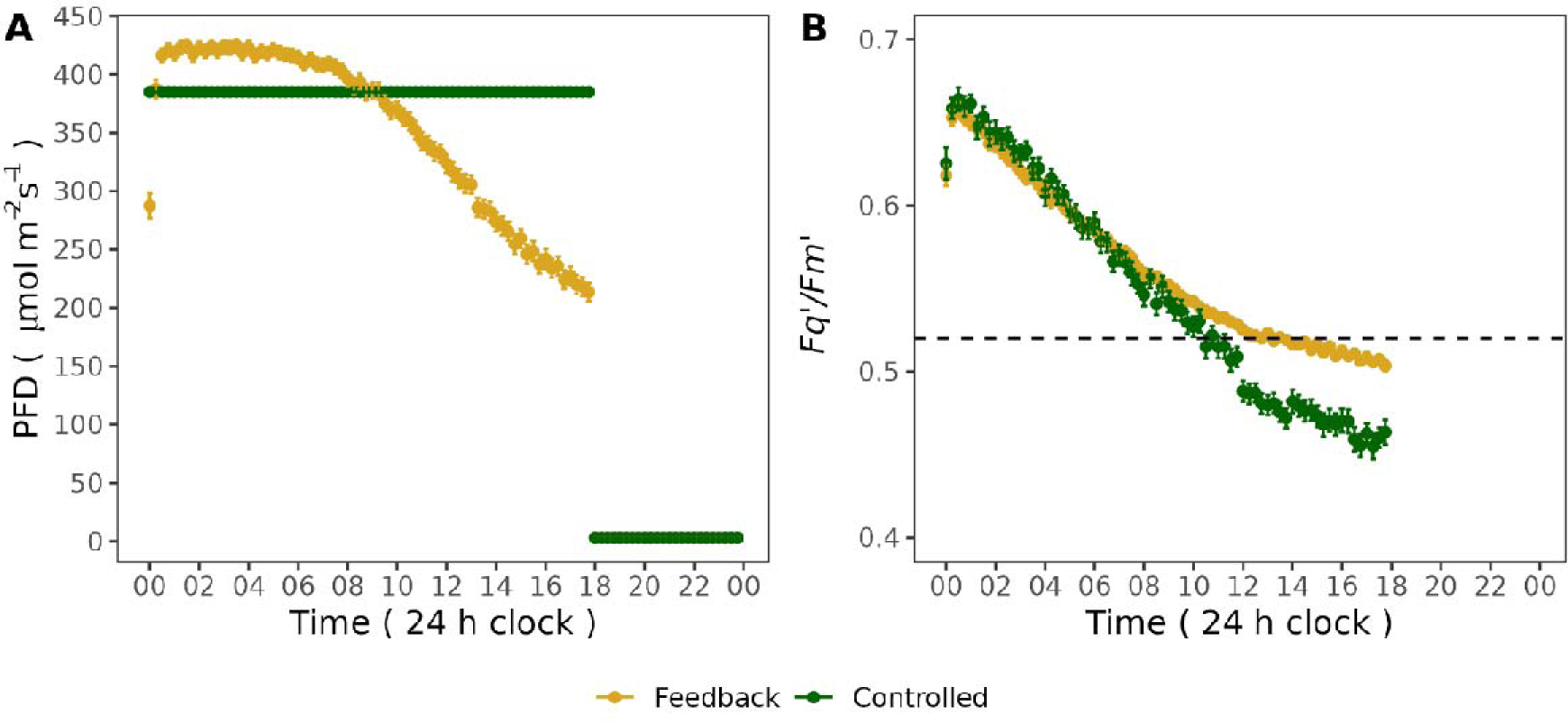
Comparison of light intensity demand and chlorophyll fluorescence between a Controlled, constant light setting (Lincoln green) and the ’Feedback’ condition of variable light setting (golden brown) for basil plants. A) While the light intensity delivered in the Controlled condition was fixed at 380 µmol m^-2^ s^-1^ for 18 h and zero for the rest of the 24 h day, in the Feedback condition, light intensity was allowed to vary between 100 and 450 µmol m^-2^ s^-1^ based on the functioning of the feedback loop with constants used as determined in Figure 3. A general pattern was established in the Feedback case of higher demand than the Controlled condition in the first ∼9 h of the day, with falling demand below that of the Controlled condition in the second half of the photoperiod. Means shown +/- SE n=6. B). No differences between treatments were observed in Fq’/Fm’ for the first ∼9 h of the day (The Feedback condition was limited to 450 µmol m^-2^ s^-1^ PFD, preventing the value of Fq’/Fm’ being forced down to the setpoint by higher light intensity). Thereafter Fq’/Fm’ observed in the Feedback condition were consistently higher than those observed for the Controlled condition, and they remained close to the setpoint for the remaining of the photoperiod. During this second phase, the minimum value of PFD of 100 µmol m^-2^ s^-1^ and limit the integral error (integral wind-up) in the first period prevented the system from continually hitting the setpoint. Means shown +/- SE , n=6.

Interestingly, while there was a 39.0 µmol m^-2^ s^-1^ difference in demanded light intensity between control and feedback at the start of the photoperiod, this was not reflected in differences in Fq’/Fm’. Overall, the photo-feedback system reduced the total light input by 2.78 mol m^-2^ d^-1^ compared to the control (p<0.01), saving 11.2% photons, along with the positive impact on growth (Figs. 4 & 5).

## DISCUSSION

Here we provide a strategy that substantially improves crop yields by 7%. In addition, this was achieved with a lower carbon footprint of 201 g CO_2_ g^-1^ FW and with energy consumption down by 6.2%. Given global food security challenges and legally binding net zero commitments, pressure has been placed on farmers and growers to deliver solutions. The technology in this study provides a straightforward, low-cost option to address these dual challenges.

In controlled environment agriculture, one of the greatest headwinds is the high energy cost of operating LED lighting^17^. Advances in LED technology and improvements in light recipes mitigate costs to some extent ^21,22, 23, 44^, although current approaches have yet to fully realise the capabilities and flexibilities of modern lighting systems. This lack of exploitation of LED technology has slowed progress towards net zero emission targets in CEA including VFs^17^. Our integrated, ‘intelligent’ systems approach (Fig. 2) can interpret real-time plant signals. We combine the latest physiological tools, modern LED technology and knowledge of plant primary metabolism to deliver adaptable and optimised lighting recipes determined by crop demand.

Crop productivity is a function of accumulated photosynthetic primary product, which is dependent on light intensity and day length. Under long-day lighting conditions (> 6 h), there is a progressive loss of photosynthetic carbon gain towards the latter part of the diurnal period. This phenomenon has been demonstrated in both field under dynamic, fluctuating conditions and even in controlled environments under constant square-wave light regimes^26,27^ (Fig. 1). The higher light level delivered to our feedback plants at the same Fq’/Fm’ level as the control suggests that further energy savings could be made by decreasing the upper limit for light intensity, without affecting productivity.

The decline in assimilation rate has been attributed to the accumulation of photosynthate^45^ and limited sink capacity^30,46^. Maintaining high light intensities when photosynthetic demand is low not only wastes energy but may also damage the plant^47,48^.

Potentially, the interaction between yield and energy use is important as growers can choose which combination of strategic outcomes they prefer. Control parameters such as the Fq’/Fm’ setpoint, maximum / minimum PFD values, and control loop constants can be manipulated to prioritise any combination of yield and energy use. For example, targeting a lower Fq’/Fm’ while increasing maximum allowable PFD when energy prices are low could deliver higher yield and shorter cycles for a constant carbon footprint. Alternatively, by increasing the Fq’/Fm’ setpoint, yield could be maintained in longer cycles with lower energy input and a reduced carbon footprint. A further capability of the system would be flexibility within cycles to adjust harvest timing dependent on the growers’ or buyers’ requirements. Crucially, the observed increases in biomass were as a consequence of photosynthetic carbon accumulation as opposed to higher leaf water content, implying no change in crop quality despite the lower energy requirement. In addition, leaves were no thinner even though leaf area was increased, accelerating carbon capture the longer a crop stays on the system.

The work covered in this paper targets outcomes based on continuous dynamic control of light intensity, yet any perturbations in plant metabolism will directly impact Fq’/Fm’^29^. Therefore, the methods outlined here can provide a sensitive indicator of plant health and stress. Photo-feedback could be used to control temperature, VPD, and water or nutrient availability in the same way as light to optimise inputs relative to outputs. Although physiological changes resulting from varying these environmental parameters often emerge over timescales longer than near-instantaneous light responses, the applicability of our approach remains. Yields are driven by both abiotic (environmental) factors as well as biotic challenges. An additional use case for our photo-feedback system is early warning of declining crop health, including, for example, disease^33,49,50^. Interventions could include application of specific spectra such as UV, or reduction in relative humidity / temperature to manage the spread of pathogens^51,52^.

Extending the control function to the broader suite of fluorescence signals beyond Fq’ / Fm’ could unlock additional applications. Using environmental measurements as inputs to a machine learning algorithm could predict the pathway to deliver fluorescence-based control solutions. Simultaneous adjustment of environmental inputs would provide the minimum-energy solution that maximised performance and yield.

Illumination is pre-eminent as a driver of productivity and has been the subject study since at least the 19^th^ Century^53^. Recent advances in LED technology have propelled academics, horticulturalists and light manufacturers to develop novel lighting recipes for yield, quality and cost. This reductionist approach has proven to be slow to translate to real-world impact given the complexity of interactions in lighting regimes. Our photo-feedback system offers a novel method for rapid prototyping and testing of candidate strategies. Iteratively assessing performance in real time could accelerate convergence toward optimal environmental settings.

While this work considered *O. basilicum* , similar patterns of diurnal photosynthetic responses have been reported for other species^42^. Chlorophyll fluorescence is a universal indicator of performance across all photosynthetic organisms in multiple growing environments^30^. The photo-feedback technology can be applied across any agricultural system where input(s) can be manipulated. Those inputs could range from pesticide application through environmental control to cultivar selection. A key feature of the system described above is that the technology is inexpensive, robust, scientifically established, and easy to use for the non-specialist. Our approach directly addresses current global challenges in food security, net zero and environmental sustainability.

## Methods

### Plants growth conditions

Seeds of *Ocimum basilicum* (sweet basil) (Suttons Seeds, Paignton, UK.), were sown in 0.29 dm^3^ plastic pots containing compost (F2S, ICL Agriculture Ltd, UK.). Pots were placed directly into the controlled environment growth chamber (Fitaclima PLH, Aralabs SA, Albarraque, Portugal), under 300 µmol m^-2^ s^-1^ Photon Flux Density (PFD) at pot height, provided by LED growth lights (Dyna, Heliospectra AB, Gothenburg, Sweden). Photoperiod was 16 h, with an air temperature of 23°C and a vapour pressure deficit (VPD) of 1.0 (±0.2) kPa. At 10 days post germination, plants were thinned to leave one individual per pot. At 14 days post germination, 24 plants were selected at random, split into control and treatments groups, and moved to a second growth environment, comprising of two independent lighting regimes. The first ‘controlled’ regime, consisted of a constant light intensity of 380 µmol m^-2^ s^-1^ at the top of the crop canopy for a 16h photoperiod. Air temperature and VPD were controlled to 22.1°C (±0.9°C) by day / 22.0°C (±0.5°C) by night and 1.07 (±0.2) kPa by day / 0.8 (±0.1) kPa by night respectively. In the second or ‘feedback’ regime, light intensity was dependent on feedback generated by a control loop based on the photophysiology of the crop.

### Control loop setup

A chlorophyll fluorimeter (PSP, ADC BioScientific, Hoddesdon, Herts, UK) measured the quantum efficiency of Photosystem II (Fq’/Fm’) every 5 minutes, in the youngest fully expanded leaf of the crop, at a constant distance (270 mm) from the growth light (Fig. 2). The fluorescence ratio measured by the fluorimeter was output as an analogue (0 to 5 v) signal and passed to an Arduino microprocessor (Uno, Arduino, Italy). A Proportional Integral (PI) feedback loop was used to compare the measured fluorescence against a setpoint intended to generate constant photosynthetic

efficiency. A PI controller seeks to minimise the error term e_(t)_ in the functions:

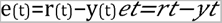

Where *e*_t_ is the error term at time *t*, expressed as the difference between the setpoint *r*_t_ (the desired value for Fq’/Fm’) and the measured value of Fq’/Fm’, *y*_t_ at that time. Having established *e*_t_, *u*_t_ is calculated:

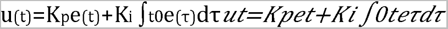

Where *u*_t_ is the new setting for the Heliospectra luminaire. The constants *K*_P_ and *K*_I_ are the multipliers for the proportional and integral elements of the control loop respectively and *e*_(τ)_ represents the errors accumulated from time *t*_0_ to present. *K*_p_ and *K*_i_ were empirically determined from observation of system performance, with values of 350 and 50 respectively selected to ensure convergence towards the setpoint (Supplementary data, dotted line) without excessive noise (e.g. Supplementary data, panel *K*_p_ = ‘1000’). No derivative was used in the process control as overshoot in the system was limited. As a result, the control loop was able to adjust light intensity in response to changes in plant photosynthetic efficiency as measured by the chlorophyll fluorescence ratio, *Fq’/Fm’*. The error between measured and setpoint values was used to calculate the output *u*_(t)_ which was then fed forward as a new setting for the Heliospectra. It was important to prevent large accumulation of errors in the integral term given the maximum and minimum light settings, which would severely affect system sensitivity. Consequently, integral wind-up was limited to the value reached when either Proportion<0 and Integral<0 and *u*_(t)_=18 or Proportion>0 and Integral>0 and *u*_(t)_=83.

All channels (representing different peak wavelength outputs of the LEDs) were given the same numeric setting in the Heliospectra software, however, light intensity outputs varied due differences in LED efficiency. The Arduino was connected to a laptop that ran a Python script communicating with the Dyna luminaire over a direct ethernet connection (Fig. 2).

The set target *Fq’/Fm’* value of 0.52 was derived from the response of *O. basilicum* photosynthetic CO_2_ uptake in parallel with changes in *Fq’/Fm’* in response to light intensity (Fig. 3A & B) using an infra-red gas analyser with an integral chlorophyll fluorimeter (Li6800, LiCor Inc., Lincoln, Nebraska, USA). This specific *Fq’/Fm’* value was close to the inflection point of the response of CO_2_ assimilation to light intensity (below the saturation point; Fig. 3A) and also between *Fq’/Fm’* values observed at the light intensity minimum and maximum PFD boundaries (Fig. 3B).

For a *Fq’/Fm’* value greater than 0.52, *e*_(t)_ was positive (i.e. plant photosynthetic efficiency was too low), tending to increase light intensity, whereas, for a value less than 0.52, *e*_(t)_ was negative and light intensity tended to decrease. To ensure that light intensities within this study were within the capability of modern LED grow lamps and therefore relevant to current indoor agricultural infrastructure, the maximum and minimum PFD values of the feedback lamp were restricted to 450 µmol m^-2^ s^-1^ and 100 µmol m^-2^ s^-1^ respectively.

### Experimental procedure

Every 24 hours before the start of the photoperiod, plants were re-randomised within light treatments. Tray height was adjusted to maintain a constant distance from the lamp to the top of the canopy, controlling for the risk of plant height systematically influencing total light capture. The plant being measured for *Fq’/Fm’*was changed at random every day. The crop was watered daily with equal quantities of water for each treatment, with an addition of nutrients (Hoaglands & Arnon, 1938) at 21 days post germination. The power consumption for each lamp was recorded using a power meter (MP001186, MultiComp, London, UK).

### Data collection and statistical analysis

After 28 days growth, leaf area, fresh and dry mass were assessed for each individual plant. Leaf area was determined using a leaf area meter (LI-3100C, Li-Cor Inc., Lincoln, Nebraska, USA). Above ground biomass (leaves and stems) were dried to constant mass at 60°C, and total dry mass measured using a 1 mg resolution balance (2006 MP, Sartorius, Germany).

Data were analysed using R statistical software in the RStudio online application^54,55^. Where possible, data were analysed as linear models, with residuals tested for normality and heteroskedasticity. For biomass and power-related data, linear mixed-effect models were used^56^, with *trial number* considered a random effect to account for the variability in the growth environment and uncertainty over the size of the plants at the start of each trial. The mixed-effect models were compared to equivalent linear models in ANOVAs ^57^ to check for best fit using the Aikike Information Criterion.

Treatment effects within the mixed-effect models were also tested as ANOVAs^57^. Effect sizes for biomass and power parameters were calculated as Cohen’s D measure.

---

**Figure S1.**
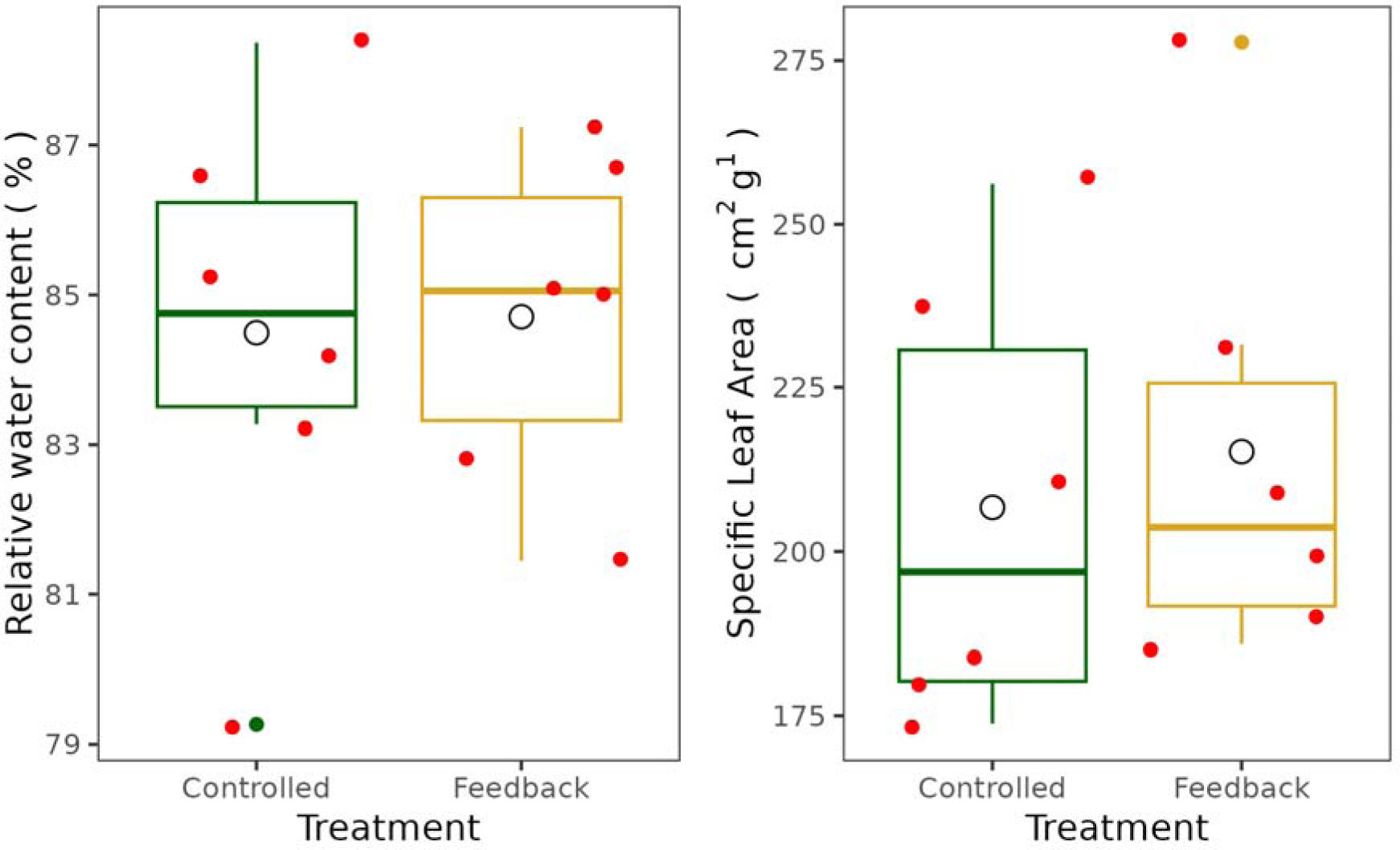
No differences in key measures of plant physiology were observed, despite the higher harvests for the Feedback treatment (see Fig. 4, main text). Data were analysed using a linear mixed effects model of the form lme(Harvest ∼ Treatment + (1|Experiment_number)). This approach captured more of the uncertainty in the (semi-controlled) environmental conditions experienced compared to a traditional linear model. Aikike Information Criteria were consistent for all results, with the mixed-effects model outperforming the linear model. A. Relative water content was the same (p=0.71) for both Controlled and Feedback regimes, B. Specific leaf area (a proxy for leaf thickness) was also the same between treatments (p=0.24). Red points represent individual experiments, black points are outliers and the black circle is the overall mean for each treatment, n=6.

**Figure S2.**
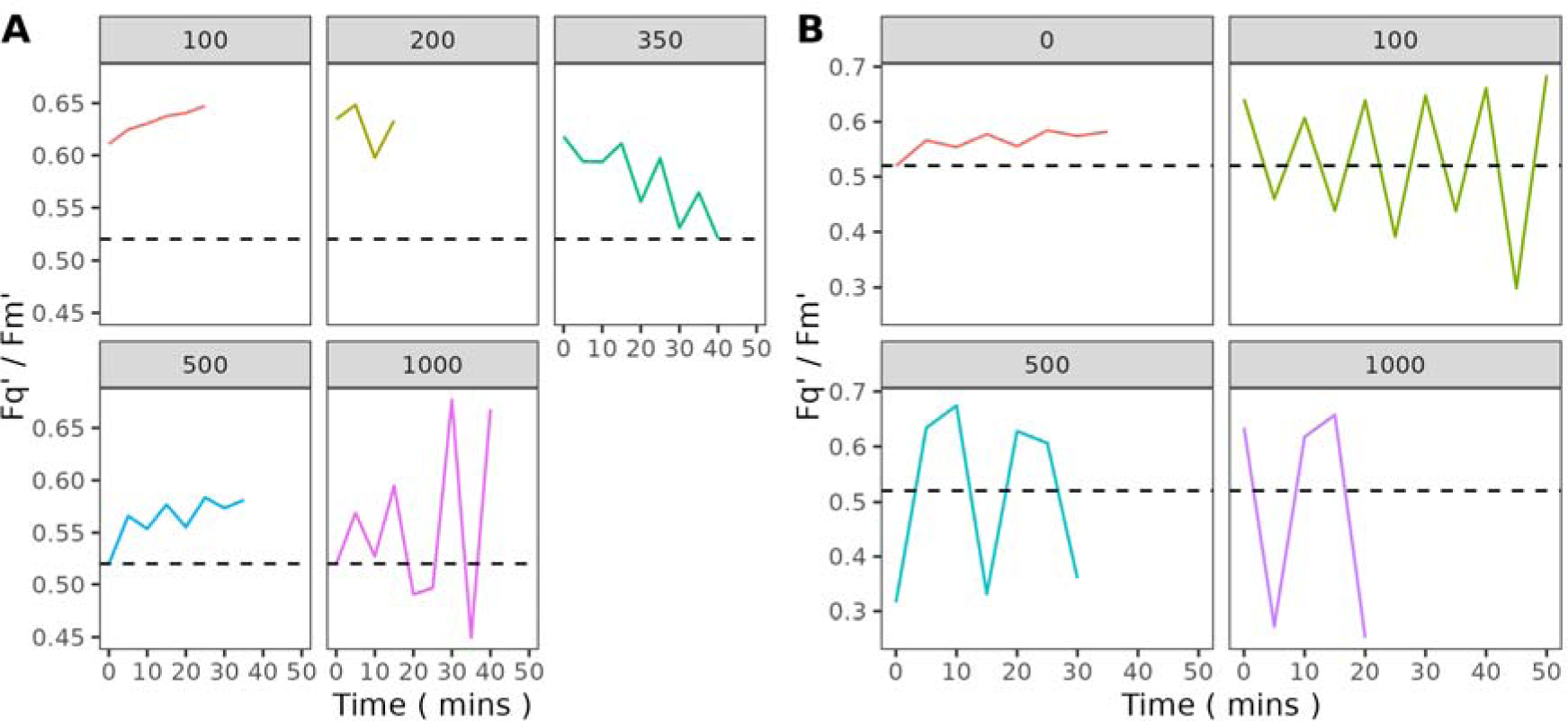
Selection of constants for the Proportion-Integral Control Loop. A. Values for the proportion constant, k_p_ were determined empirically, with each panel representing a time series of observations at a given constant. Values that rapidly returned Fq’/Fm’ to the setpoint without overshooting were preferred. As a result a value of 350 was selected. B. In a similar manner, the value for the integral constant in the control loop was selected empirically. Here, K_i_ = 50 was selected, lying between the K = 0 and K = 100 results shown.

## References

1. Tilman, D., Balzer, C., Hill, J., & Befort, B. L. Global food demand and the sustainable intensification of agriculture. Proceedings of the national academy of sciences 108(50), 20260–2026 (2011).

2. Foley, J. A. Can we feed the world sustain the planet? Scientific American 305(5), 60–65 (2011).

3. Foley, J.A., Ramankutty, N., Brauman, K.A., Cassidy, E.S., Gerber, J.S., Johnston, M., Mueller, N.D., O’Connell, C., Ray, D.K., West, P.C., Balzer, C., Bennett, E.M., Carpenter, S.R., Hill, J., Monfreda, C., Polasky, S., Rockstrom, J., Sheehan, J., Siebert, S., Tilman, D. & Zaks, D.P.M. Solutions for a cultivated planet. Nature 478, 337–342 (2011)

4. FAO. Greenhouse gas emissions from agrifood systems. Global, regional and country trends, 2000-2020. *FAOSTAT Analytical Brief Series*50, 1-12 (2022).

5. Ray, D. K., Mueller, N. D., West, P. C., & Foley, J. A. Yield trends are insufficient to double global crop production by 2050. PloS one, 8(6), e66428 (2013).

6. Shamshiri, R.R., Kalantari, F., Ting, K.C., Thorp, K.R., Hameed, I.A., Weltzien, C., Ahmad, D. & Shad, Z. Advances in greenhouse automation and controlled environment agriculture: A transition to plant factories and urban agriculture. International Journal of Agricultural and Biological Engineering, 1–22 (2018).

7. Gruda, N., & Tanny, J. Protected crops. Horticulture: Plants for People and Places, Volume 1: Production Horticulture, 327–405 (2014).

8. Gruda, N. & Tanny, J. Protected crops – recent advances, innovative technologies and future challenges. Acta Hortic. 1107, 271–278 (2015).

9. Van Gerrewey, T., Boon, N., & Geelen, D. Vertical farming: The only way is up? Agronomy 12(1), 2 (2021).

10. van Iersel, M.W. & Gianino, D. An adaptive control approach for light-emitting diode lights can reduce the energy costs of supplemental lighting in greenhouses. Horticultural Science 52, 72–77 (2017).

11. Wong, C., Wood, J. & Paturi, S. Vertical farming: an assessment of Singapore City. Etropic: electronic journal of studies in the tropics 19, 228–248 (2020).

12. Van Delden, S.H., SharathKumar, M., Butturini, M., Graamans, L.J.A., Heuvelink, E., Kacira, M., Kaiser, E., Klamer, R.S., Klerkx, L., Kootstra, G. and Loeber, A. Current status and future challenges in implementing and upscaling vertical farming systems. Nature Food 2(12), 944–956 (2021).

13. Beacham, A. M., Vickers, L. H., & Monaghan, J. M. Vertical farming: a summary of approaches to growing skywards. The Journal of Horticultural Science and Biotechnology 94(3), 277–283 (2019).

14. Bian, Z., Jiang, N., Grundy, S., & Lu, C. Uncovering LED light effects on plant growth: New angles and perspectives-LED light for improving plant growth, nutrition and energy-use efficiency. In *International Symposium on New Technologies for Environment Control*, Energy-Saving and Crop Production in Greenhouse and Plant 1227, 491–498 (2017).

15. Abdullah, M.J., Zhang, Z. & Matsubae, K. Potential for Food Self-Sufficiency Improvements through Indoor and Vertical Farming in the Gulf Cooperation Council: Challenges and Opportunities from the Case of Kuwait. Sustainability 13, 12553 (2021).

16. Graamans, L., van den Dobbelsteen, A., Meinen, E., & Stanghellini, C. Plant factories; crop transpiration and energy balance. Agricultural Systems 153, 138–147 (2017).

17. Stanghellini, C., & Katzin, D. (2024). The dark side of lighting: a critical analysis of vertical farms’ environmental impact. Journal of Cleaner Production , 142359.

18. Casey, L., Freeman, B., Francis, K., Brychkova, G., McKeown, P., Spillane, C., Bezrukov, A., Zaworotko, M. and Styles, D. Comparative environmental footprints of lettuce supplied by hydroponic controlled-environment agriculture and field-based supply chains. Journal of Cleaner Production 369, 133214 (2022).

19. Cowan, N., Ferrier, L., Spears, B., Drewer, J., Reay, D., & Skiba, U. CEA systems: the means to achieve future food security and environmental sustainability? Frontiers in Sustainable Food Systems 6, 891256 (2022).

20. Palmer, S. & van Iersel, M.W. Increasing growth of lettuce and mizuna under sole-source LED lighting using longer photoperiods with the same daily light integral. Agronomy 10, 1659 (2020).

21. Nelson, J. A., & Bugbee, B. Economic analysis of greenhouse lighting: light emitting diodes vs. high intensity discharge fixtures. PloS one 9(6), e99010 (2014).

22. Bugbee, B. Economics of LED Lighting. In: Dutta Gupta, S. (eds) Light Emitting Diodes for Agriculture (Springer, 2017).

22. van Iersel, M. W. Optimizing LED lighting in controlled environment agriculture. Light emitting diodes for agriculture: Smart lighting, 59–80 (Springer, 2017).

24. Austin, R. B. Crop characteristics and the potential yield of wheat. The Journal of Agricultural Science 98(2), 447–453 (1982).

25. Matthews, J. S., Vialet-Chabrand, S. R., & Lawson, T. Diurnal variation in gas exchange: the balance between carbon fixation and water loss. Plant Physiology 174(2), 614–623 (2017).

26. Matthews, J. S., Vialet-Chabrand, S., & Lawson, T. Acclimation to fluctuating light impacts the rapidity of response and diurnal rhythm of stomatal conductance. Plant Physiology 176(3), 1939–1951 (2018).

27. Vialet-Chabrand, S., Matthews, J. S., Simkin, A. J., Raines, C. A., & Lawson, T. Importance of fluctuations in light on plant photosynthetic acclimation. Plant physiology 173 (4), 2163–2179 (2017).

28. Salter, W. T., Gilbert, M. E., & Buckley, T. N. A multiplexed gas exchange system for increased throughput of photosynthetic capacity measurements. Plant Methods 14, 1–12 (2018).

29. Baker, N. R. Chlorophyll fluorescence: a probe of photosynthesis in vivo. Annual Review of Plant Biology 59, 89–113 (2008).

30. Murchie, E. H., & Lawson, T. Chlorophyll fluorescence analysis: a guide to good practice and understanding some new applications. Journal of experimental botany 64(13), 3983–3998 (2013).

31. Simkin, A. J., López-Calcagno, P. E., & Raines, C. A. Feeding the world: improving photosynthetic efficiency for sustainable crop production. Journal of Experimental Botany 70(4), 1119–1140 (2019).

32. Cavender-Bares, J., & A. Bazzaz, F. From leaves to ecosystems: using chlorophyll fluorescence to assess photosynthesis and plant function in ecological studies. In Chlorophyll a fluorescence: a signature of photosynthesis , 737–755 (Springer, 2004).

33. Gorbe, E., & Calatayud, A. Applications of chlorophyll fluorescence imaging technique in horticultural research: A review. Scientia Horticulturae 138, 24–35 (2012).

34. Elkins, C., & van Iersel, M. W. Longer photoperiods with the same daily light integral increase daily electron transport through photosystem II in lettuce. Plants 9(9), 1172 (2020a).

35. Elkins, C., & van Iersel, M. W. Longer photoperiods with the same daily light integral improve growth of rudbeckia seedlings in a greenhouse. Horticultural Science 55(10), 1676–1682 (2020b).

36. Palmer, S. & van Iersel, M.W. Increasing growth of lettuce and mizuna under sole-source LED lighting using longer photoperiods with the same daily light integral. Agronomy 10, 1659 (2020).

37. Weaver, G. & van Iersel, M.W. Longer photopheriods with adaptive lighting control can improve growth of greenhouse-grown ‘Little Gem’ lettuce (Latuca sativa). Horticultural Science 55, 573–580 (2020).

38. van Iersel, M.W., Mattos, E., Weaver, G., Ferrarezi, R.S., Martin, M.T., Haidekker, M. Using chlorophyll fluorescence to control lighting in controlled environment agriculture. Acta Horticulturae 1134, 427–434 (2016a)

39. van Iersel, M.W., Weaver, G., Martin, M.T., Ferrarezi, R.S., Mattos, E., Haidekker, M. A chlorophyll fluorescence-based biofeedback system to control photosynthetic lighting in controlled environment agriculture. Journal of the American Society for Horticutural Science. 141(2), 169–176 (2016b).

40. Baker, N. R., & Rosenqvist, E. Applications of chlorophyll fluorescence can improve crop production strategies: an examination of future possibilities. Journal of experimental botany 55(403), 1607–1621 (2004).

41. Pocock, T. Light-emitting diodes and the modulation of specialty crops: light sensing and signaling networks in plants. Horticultural Science 50, 1281–1284 (2015).

42. Suwannarut, W., Vialet-Chabrand, S., & Kaiser, E. Diurnal decline in photosynthesis and stomatal conductance in several tropical species. Frontiers in Plant Science 14, 1273802 (2023).

43. Kaiser, E., Morales, A., Harbinson, J., Kromdijk, J., Heuvelink, E., & Marcelis, L. F. Dynamic photosynthesis in different environmental conditions. Journal of Experimental Botany 66(9), 2415–2426 (2015).

44. Kusuma, P., Pattison, P.M. and Bugbee, B. From physics to fixtures to food: Current and potential LED efficacy. Horticulture research 7, 56 (2020).

45. Stitt, M. Rising CO2 levels and their potential significance for carbon flow in photosynthetic cells. Plant, Cell & Environment 14(8), 741–762 (1991).

46. Matthew, J.P. & Foyer, C.H. Sink regulation of photosynthesis, Journal of Experimental Botany 52 (360), 1383–1400 (2001).

47. Powles, S. B. Photoinhibition of photosynthesis induced by visible light. Annual review of plant physiology 35(1), 15–44 (1984).

48. Raven, J. A. The cost of photoinhibition. Physiologia plantarum 142(1), 87–104 (2011).

49. Scholes, J. D., & Rolfe, S. A.Chlorophyll fluorescence imaging as tool for understanding the impact of fungal diseases on plant performance: a phenomics perspective. Functional Plant Biology 36(11), 880–892 (2009).

50. Wang, H., Qian, X., Zhang, L., Xu, S., Li, H., Xia, X., Dai, L.,Xu, L., Yu, J. & Liu, X. A method of high throughput monitoring crop physiology using chlorophyll fluorescence and multispectral imaging.Frontiers in Plant Science 9, 407 (2018).

51. Elad, Y., & Pertot, I. Climate change impacts on plant pathogens and plant diseases. Journal of Crop Improvement 28(1), 99–139 (2014).

52. Kunz, B. A., Cahill, D. M., Mohr, P. G., Osmond, M. J., & Vonarx, E. J. Plant responses to UV radiation and links to pathogen resistance. International Review of Cytology 255, 1–40 (2006).

53. Darwin, C.R. On the movements and habits of climbing plants, Journal of the Linnean Society of London (Botany) 9, 1–118 (1865).

54. 54. R Core Team. R: A language and environment for statistical computing. R Foundation for Statistical Computing, at https://www.R-project.org/ (2021).

55. 55. RStudio Team. RStudio: Integrated Development for R. RStudio, PBC at RL http://www.rstudio.com/ (2020).

56. Bates, D., Mächler, M., Bolker, B., & Walker, S. Fitting linear mixed-effects models using lme4. Journal of Statistical Software 67(1), 1–48 (2015).

57. Fox, J., & Weisberg, S. An R companion to applied regression (Sage publications, 2018)

