## Supplementary material for "Green Conversations: Harnessing Plant Communication to control growth light intensity": Suppl: Supplementary Figure Legends.pdf

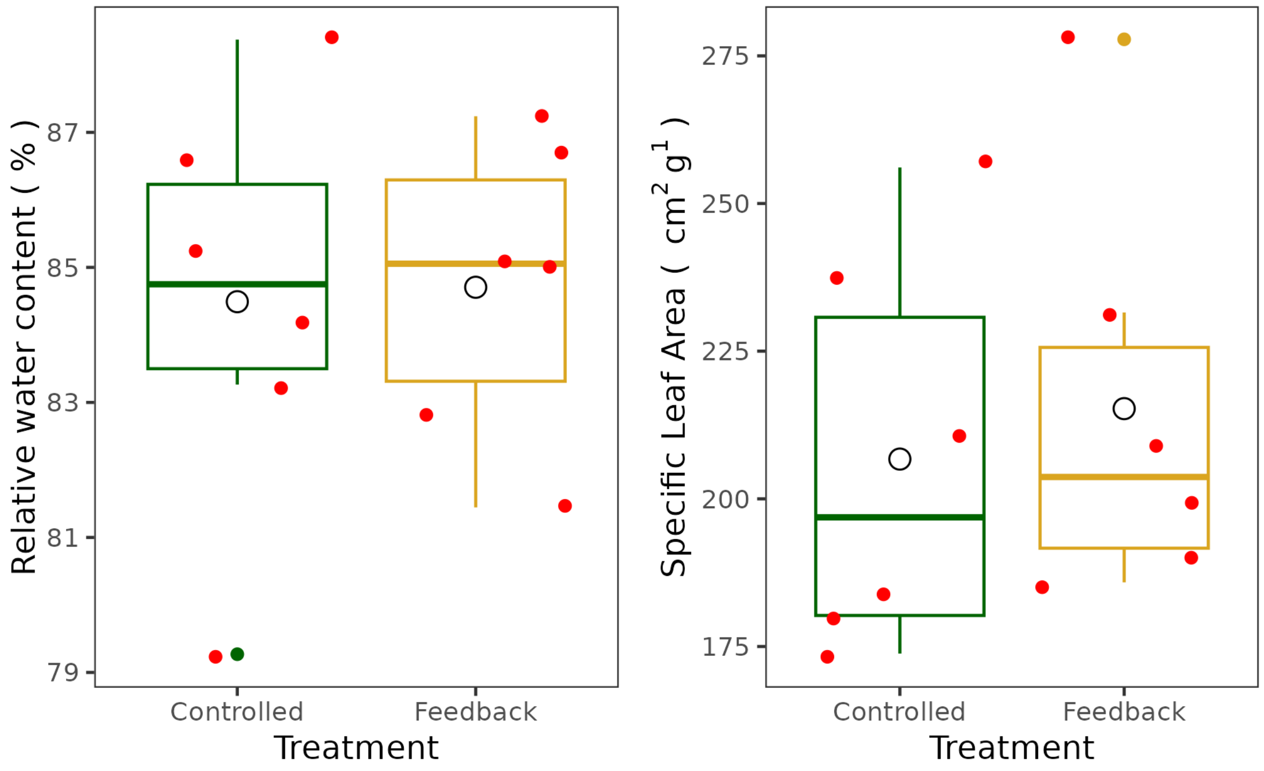


Figure S1. No differences in key measures of plant physiology were observed, despite the higher harvests for the Feedback treatment (see Fig. 4, main text). Data were analysed using a linear mixed effects model of the form lme(Harvest ~ Treatment + (1|Experiment_number)). This approach captured more of the uncertainty in the (semi-controlled) environmental conditions experienced compared to a traditional linear model. Aikike Information Criteria were consistent for all results, with the mixed-effects model outperforming the linear model. A. Relative water content was the same ($p=0.71$) for both Controlled and Feedback regimes, B. Specific leaf area (a proxy for leaf thickness) was also the same between treatments ($p=0.24$). Red points represent individual experiments, black points are outliers and the black circle is the overall mean for each treatment, $n=6$.

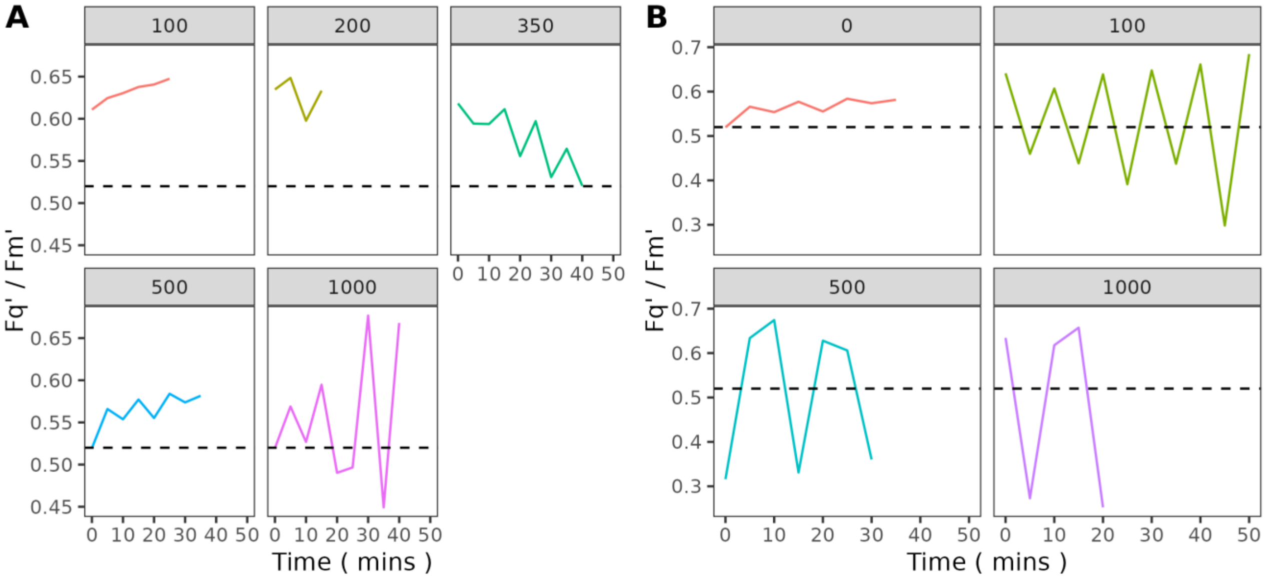


Figure S2. Selection of constants for the Proportion-Integral Control Loop. A. Values for the proportion constant, $k_p$ were determined empirically, with each panel representing a time series of observations at a given constant. Values that rapidly returned *Fq'/Fm'* to the setpoint without overshooting were preferred. As a result a value of 350 was selected. B. In a similar manner, the value for the integral constant in the control loop was selected empirically. Here, $K_i$ = 50 was selected, lying between the K = 0 and K = 100 results shown.
